# Computing Minimal Boolean Models of Gene Regulatory Networks

**DOI:** 10.1101/2021.05.22.445266

**Authors:** Guy Karlebach, Peter N Robinson

## Abstract

Models of Gene Regulatory Networks (GRNs) capture the dynamics of the regulatory processes that occur within the cell as a means to understand the variability observed in gene expression between different conditions. Arguably the simplest mathematical construct used for modeling is the Boolean network, which dictates a set of logical rules for transition between states described as Boolean vectors. Due to the complexity of gene regulation and the limitations of experimental technologies, in most cases knowledge about regulatory interactions and Boolean states is partial. In addition, the logical rules themselves are not known a-priori. Our goal in this work is to create an algorithm that finds the network that fits the data optimally, and identify the network states that correspond to the noise-free data. We present a novel methodology for integrating experimental data and performing a search for the optimal consistent structure via optimization of a linear objective function under a set of linear constraints. In addition, we extend our methodology into a heuristic that alleviates the computational complexity of the problem for datasets that are generated by single-cell RNA-Sequencing(scRNA-Seq). We demonstrate the effectiveness of these tools using a public scRNA-Seq dataset and the GRN that is associated with it. Our methodology will enable researchers to obtain a better understanding of the dynamics of gene regulatory networks and their biological role.

## 1. Introduction

Maintenance of cellular functions requires the orchestration of many interleaving processes over time and space. A Gene Regulatory Network (GRN) is a set of genes such that the present state of their expression trajectory can be predicted from past states via the regulatory relationships between the genes. As a model that can display complex behavior and at the same time is straightforward to specify, GRNs have been used to describe the regulation of process as different as cell differentiation, circadian clocks and diauxic shift [3], [11], [18]. Consequently, many methods for reconstructing GRNs from experimental data at varying levels of detail have been proposed [5], [7], [17]. Arguably the most basic formulation, the Boolean network, describes gene expression states as Boolean values and changes in those levels as Boolean functions [9]. While the simplicity of this model imposes a certain level of abstraction, it also makes it applicable to a broader range of datasets and simplifies its analysis. Interestingly, despite the relative simplicity of Boolean networks, fitting a Boolean network to a gene expression dataset given a set of regulatory relationships is an NP-Hard problem [8]. In practice the regulatory interactions that occurs in a given dataset are not known, which can result in redundant regulators after fitting. Not surprisingly, an algorithm for optimal fitting that takes into account the complexity of network structure has not been proposed to-date [1], [12], [15]. In this paper we present a novel algorithm for finding the optimal Boolean network structure with respect to its fit to a dataset, and for denoising the data. We show that minimizing the encoding of the noise and the network structure can be formulated as a 0/1 Integer Linear Programming problem (ILP), and thus solved using powerful branch and bound methods that exist for ILP. In addition, we provide heuristic that alleviates the computational complexity of the problem, and can be used to solve the problem or as a subroutine for finding bounds for the 0/1 ILP solution. We demonstrate the effectivenes of our methodology on a gene expression dataset of single-cell RNA Sequencing from mouse embryonic midbrain.

## 2. Methods

### Optimal Fit of Boolean Networks

A gene expression dataset consists of a N × M matrix, where N corresponds to the number of genes whose expression level was measured, and M corresponds to the number of experiments. The expression values in the matrix can be discrete or continuous, depending on the experimental technology that was used for generating the data, and in both cases captures the variation in transcriptional regulation and contains noise. In order to map these values to Boolean values one needs to label each observation as belonging to a state of low or high expression. The mapping does not need to be perfect, since any analysis method should be able to account for some degree of incorrect mappings as a result of noise. As the methodology proposed in this section is independent of the choice of a mapping, in the rest of this section we will assume that the mapping has already been applied to the data. In the Results section we demonstrate the process using an experimental dataset.

A trajectory of a Boolean network is a sequence of states such that each state except for the first state in the sequence is derived from the previous state using a set of Boolean functions, also known as logic tables. Each Boolean function determines the value of exactly one gene, and its inputs are the Boolean values of any number of genes in the network. Usually, it is assumed that the number of inputs is small compared to the total number of genes. The regulatory relationships of a Boolean network can be illustrated as edges from inputs to outputs in a directed graph, called a regulation graph. A gene that has an outgoing edge to another gene is referred to as a regulator, and a gene with an incoming edge as a target (a gene can be both a regulator and a target in the same network). A steady state is a state that repeats itself in a trajectory indefinitely unless perturbed by external signals, i.e. signals that are not part of the network. In a typical gene expression dataset the experiments correspond to a single time point, and therefore the network is assumed to be in a steady state in each experiment. For simplicity of description we assume in the rest of this section that the network is in steady state, however the algorithm presented here is applicable to time-series as well. This is easy to see if we convert the trajectory to an undirected regulation graph, where each data point corresponds to a node/gene, and a node/gene in time t is regulated by the nodes at time t-1 that correspond to its regulators in the original regulation graph. Then the same variables describe the logic tables of all the copies of the same gene, i.e. the nodes that correspond to it in different time points.

Discrepancies between a dataset and a network model occur when the Boolean values in an experimental dataset do not agree with any network trajectory due to experimental noise. This presents a difficulty if the network model is not known a-priori, since enumerating all possible networks is infeasible. Formally, let *g_i_* denote the integer index of the *i^th^* gene in the list of genes, let 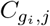 denote the Boolean value of gene *g_i_* in experiment *j*, and let 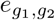 denote a directed edge between genes *g*_1_ and *g*_2_. We say that the data contains a discrepancy if for some gene index *g* and two experiments *i*_1_ and *i*_2_, 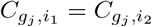 for all genes *g_j_* such that an edge 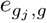 exists, but 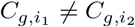. It follows from the network’s determinism that at least one of the experiments *i*_1_ or *i*_2_ does not agree with any network trajectory. To simplify the notation, we will refer to a gene by its name or index in the list of genes interchangeably. The number of regulators of a gene *g* will be denoted as *indegree*(*g*).

Assuming that *P* ≠ *NP*, there does not exist a polynomial time algorithm for resolving all the discrepancies with the minimal number of changes. Therefore, either a heuristic that finds a local optimum or an algorithm that may not terminate in a reasonable amount of time must be used instead. Another difficulty is that a strict subset of the regulation graph may provide a better fit to the data, as not all the interactions occur under every condition, and so the structure of the network itself needs to be considered in the search for the optimal solution. This adds another level of complexity to the already difficult problem. As it turns out, the problem can be formulated as 0/1 ILP, as described in the next subsection.

### An Algorithm for the Optimal Minimal Network

The in-degree of nodes in the regulation graph is usually assumed to be small compared to the number of genes or the number of experiments. If we assume that it is a constant value in terms of computational complexity, we can define a set of constraints on the values that have to be changed in order to remove all discrepancies from the data. Let *C_i,j_* denote the Boolean input value of gene *i* at experiment *j*, and let 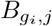 equal 1 if the value of gene *g_i_* in experiment *j* was flipped in the solution (i.e. did not fit the trajectories of the inferred network), and otherwise 0. Then for every experiment *j* and for every gene *g*_*k*+1_ with regulators *g*_1_, *g*_2_, …, *g_k_* and for every Boolean vector (*w*_1_, *w*_2_, …, *w_k_*), *w_j_* ∈ {0, 1}, if the output of the Boolean function that determines the value of *g*_*k*+1_, *I*(*w*_1_, *w*_2_, …, *w_k_*), is 1, the following constraint must hold:

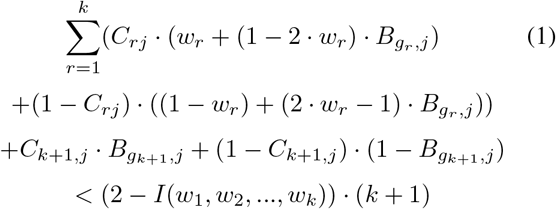

This constraint means that if the output variable *I*(*w*_1_, *w*_2_, …, *w_k_*) was set to 1, whenever the inputs *w*_1_, *w*_2_, …, *w_k_* appear in the solution, the output (the value of *g*_*k*+1_) must be 1. Similarly, if *I*(*w*_1_, *w*_2_, …, *w_k_*) is set to 0 the following constraint must hold:

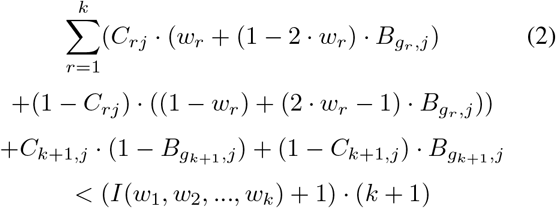

By requiring that under these constraints the following sum is minimized:

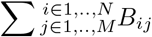

we can use a branch and bound algorithm for 0/1 integer programming to find a solution that fits the data with a minimal number of changes, and construct a new dataset with values *D_ij_* = ((*C_ij_* + *B_ij_*) mod 2), *i* ∈ 1..*N, j* ∈ 1..*M* . However, this formulation assumes that all regulatory interactions take effect in the data, which is rarely the case in practice. In order to choose the solution that also minimizes the network structure, for every gene *g_i_* and each one of its targets *g_j_*, we create another Boolean variable *R_ij_*, that is constrained to be greater equal than every difference between regulatory outputs (the *I* variables) where pairs of logic table rows are identical in all inputs except *g_i_*. If the regulatory output changes when only the regulator *g_i_* changes its value, it will be constrained to equal 1. Since other inputs can be removed as well, we add the same constraints to *R_ij_* even for pairs of rows where other inputs change, and we add the R variables of the changing inputs with a minus sign to the right hand side, i.e subtract them from the difference between the *I* variables. If the other variables that change have been removed, *R_ij_* will still be subjected to the same constraint, and will be set to 1 if the output changes. If the new constraints can be satisfied without setting *R_ij_* to 1, then the edge from *g_i_* to *g_j_* in the regulation graph is redundant, because it does not explain any change that is not explained by other edges. Using the *R_ij_* variables, we can also weigh the size of the logic tables they contribute to, by defining a new type of variable 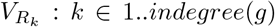 that sums the number of *R_ij_* variables of a gene *g* (the gene index is removed from the variable name for simplicity). This variable will be constrained to 1 if at least R of the variables if gene *g* are set to 1. For example, for a gene with 3 regulators, 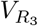 is a Boolean variable which is greater equal the sum of the three regulator variables of the gene divided by 3, minus 2/3, so it is only constrained to equal 1 if all three regulators of the gene are kept. Similarly, 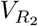 is a Boolean variable which is greater or equal any sum of two regulator variables of the gene divided by 2 minus 1/2, and 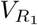 is a Boolean variable greater or equal any single regulator variable of the gene. The weights of these variables in the objective function are 2 for 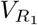, 2^2^ − 2 for 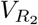 and 2^3^ − 2^2^ for 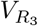. If 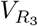 is equal 1, so do the other two variables, and therefore the objective function is added with the size of a Boolean table of 3 regulators, i.e 2^3^. Similarly, if 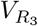 is 0 but 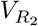 is equal 1, so is 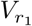, and so the addition to the objective function is 2^2^. If a single regulator is chosen, then the size of the logic table is 2 and that is the addition to the objective function. This example is trivially generalized to any number of regulators. The number of variables in the resulting 0/1 integer linear programming is 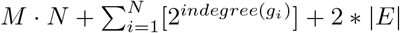. The representation of the regulation graph requires *log*_2_(*N*) bits per edge, and one bit to represent either the separator between edges or the end of the list of a given gene’s edges. A Boolean vector of length N, whose entries correspond to genes, represents which genes have regulators. Each regulation function output requires one bit. The noise representation requires a Boolean vector of length *M* · *N*, where a 0 represent the absence of noise and 1 represents noise. To minimize the edit distance from a state of ignorance, where the network is empty and noise is absent, we weigh the B variales in the objective function with 1, add 1 to the weight of the first V variable of each gene(i.e. when *k* = 1), and each R variable is weighed with *log*_2_(*N*) + 1. Minimization of the objective will then minimize the representation of the network and the noise, as desired.

### A Heuristic for Single Cell Datasets

Speeding up the solution for computationally hard problems has been an active topic of Bioinformatics research of scRNA-Seq [19], [20]. A heuristic for solving the fitting problem would enable fitting for hard instances, and could also be used for improving the upper bound during the 0/1 ILP search. Therefore, we propose the following heuristic for scRNA-Seq data:

1. Cluster the input states into K clusters.
2. Solve the problem exactly for the set of cluster centers, where fractional values in the centers are rounded to the closer bit, to obtain the set of regulators *E*, then find the de-noised states using the fixed set *E* as described in the previous section, to obtain a de-noised set of states *H*^∗^.
3. Initialize the full solution *H* to the empty set. For input state *i* ∈ 1..*M*, add it to *H*^∗^ and if it conflicts with a state in *H*^∗^ with respect to *E*, change it incrementally to match the entries of the state in *H*^∗^ that is closest to it, until all discrepancies are resolved, and then add it to the full solution *H* as well. Otherwise, if it does not conflict add it to *H* and *H*^∗^ without change.
4. Return *H*

By the construction of *H*^∗^, it is free from discrepancies. *H* is built incrementally such that after every addition of a state it is free from discrepancies, and therefore it is also free of discrepancies. It is easy to see that when step 3 is completed, either a new state that is free of discrepancies or an existing state, that by the loop invariant is free of discrepancies, is added to *H* and *H*^∗^. Similarly, step 3 preserves the set of regulators inferred in step 2, and therefore it does not increase the number of regulators. In single-cell data, clusters of cells will have a similar network state, and thus it is likely that a close state to the one that is present in the optimal solution will already be included in *H*^∗^. This will provide an upper bound for the number of changes applied to each state. Note that the optimal order by which changes are applied to the conflicting state in step 3 may depend on the input. For example, one may wish to order the changes according to the number of discrepancies that they resolve after the last change, or to choose the order based on the network structure. Similar considerations can be applied to the order by which states are selected for addition, for example, by the number of discrepancies with states that have not been added yet. The weight of a discrepancy in step 1 can be set to the number of states in its cluster, as each cluster represents a set of states that are fitted to the network’s trajectories.

## Results

### Analysis of the Gene Regulatory Network for Mid-brain Dopaminergic Neurons

In order to test our algorithm we use the mouse embryo scRNA dataset of LaManno et al. [10] and the midbrain dopaminergic (mDA) neuron developmental GRN that was described by [2]. To obtain the gene counts we used the scRNA R package [13]. The R package Seurat [14] provides function to inspect the data and determine the threshold for screening out cells with an unusually high or low number of features, leading to lower and upper thresholds of 500 and 4,000 features for this dataset, respectively. After filtering, the dataset contained 1,631 cells (experiments). Since scRNA-Seq data contains a high number of 0 counts, we applied SAVER [6] to impute expression values for the network genes. To obtain Boolean values, we map values smaller or equal to the median expression value to Boolean 0, and all other counts to 1. The number of clusters K used in the heuristic was set to 50. The clustering algorithm that we used was k-means as implemented in R, with all other parameters at their default values. For solving the 0/1 ILP problem we used Gurobi [4] The optimal solution flips 5,997 Boolean values, which corresponds to a noise level of approximately 17.5% of the input values. The observed number of discrepancies with respect to the original network structure is 8,331, approximately 24.3% of the input values.

### Minimal Network Structure and Logic

Figure 1 shows a multidimensional scaling of the samples using the binary distance between them after network fitting. Each circle corresponds to a distinct network state. The circles form several clusters which correspond to states that are close to one another in the state space. We propose that each cluster corresponds to a distinct network function, and cells in the same cluster thus share a phenotype induced by the network. Using the inferred logic, perturbations to drive the network to each of these states could be derived, and phenotypic differences under these perturbations could be further studied. Examination of the optimized network structure shows that fitting has resulted in a sparser network structure, as illustrated in Figure 2. After fitting, only 43 of the original 51 edges remained in the network, indicating that some of the interactions do not take place under the conditions of the study. Figure 3 shows the original network, where edges that were removed after fitting are highlighted in gold. In order to characterize the complexity of the regulatory logic, we examined for all genes the correlation between the number of expressed regulators and the expression of the target. The number of expressed regulators was far from perfectly correlated with the regulatory logic’s output when the genes had multiple regulators affecting its value, reflected by low absolute values for the Kendall’s tau statistic, with a median of 0.13 and a mean of 0.24. Pitx3 had the highest correlation value (0.57) among genes that had more than 2 regulators. These results suggest that the network’s logic is not a monotone function of the number of regulators, and highlights the importance of models that allow for complex regulation functions.

**Figure 1.**
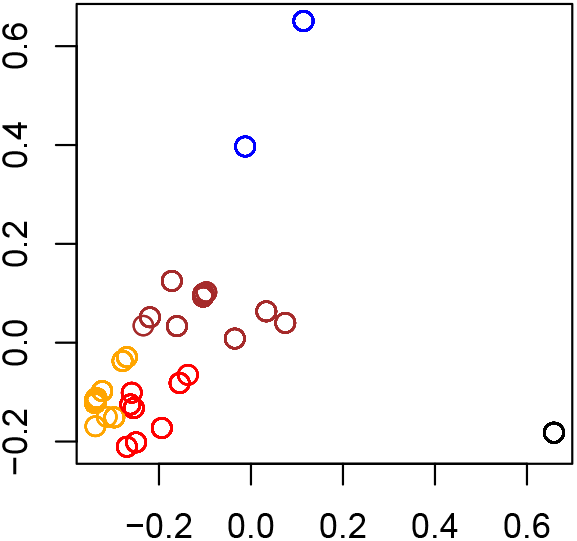
Multidimensional scaling of network states

**Figure 2.**
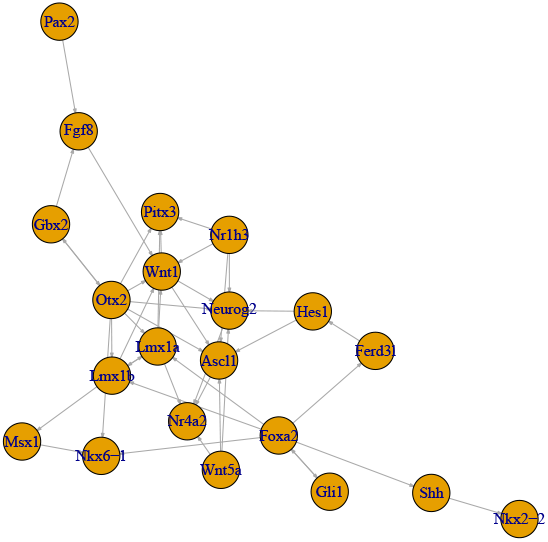
Optimized network structure

**Figure 3.**
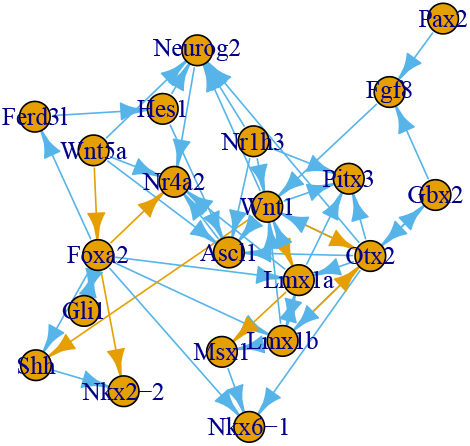
Network structure before optimization

### Noise Distribution Across Network Nodes

The noise level of different genes in the model is of interest since assumptions about this parameter are often made in models of gene expression. Therefore, we examined for each gene the number of input values of each type (Boolean 0 and 1) that were flipped in the optimal fit. The distribution of noise across the genes was non-uniform (Fig. 4, Kolmogorov-Smirnov p-value 1.2 · 10^−8^) and was not associated with an expressed state (Boolean 1) or non-expressed state(Boolean 0). This suggests that modeling assumptions that assign an equal level of noise for all genes may lead to wrong conclusions. Further research into the nature of the differences in noise levels between the different genes in this type of data could provide the basis for further improvement of modeling methods.

**Figure 4.**
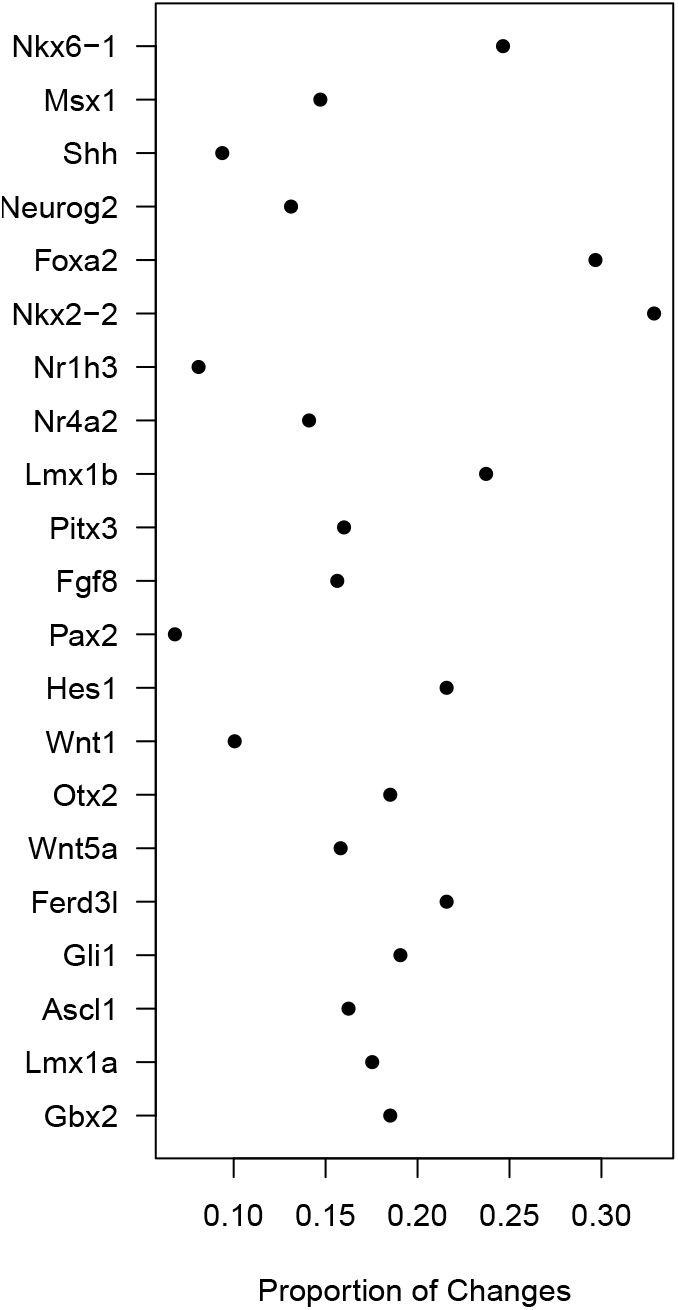
Noise distribution across genes

## Conclusion

We propose a new algorithm for fitting a Boolean network model to gene expression data that finds an optimal solution with respect to network structure and fit to the data. We further present a heurisitic that alleviates the computational complexity of the problem and therefore provides a practical solution for cases in which an exact solution cannot be obtained, which are bound to exist assuming P*neq*NP. Using known regulatory relationships and a dataset of scRNA-Seq measurements, we demonstrated our algorithm by inferring the network structure and its state in different cells. Inspection of the structural properties of the inferred network show that fitting prunes redundant regulators, and stresses the importance of sparsity constraints in the search algorithm. Furthermore, the inferred logic was diverse and showed that constraints on the logic functions should be avoided. By examining the de-noised data we found that distinct regulatory trajectories could give rise to different types of cells. Thus, the regulatory relationships between transcription factors and their targets can enable the selection of targets for perturbation experiments to validate phenotypes induced by the modeled gene regulatory network.

